# The Development and Validation of a Portable, Benchtop, Hydrostatic Model to Acquire and Analyze Quantitative Data on Unused and Failed Hydrocephalus Catheters

**DOI:** 10.1101/2022.10.04.510886

**Authors:** Pranav Gopalakrishnan, Ahmad Faryami, Carolyn A. Harris

## Abstract

**Introduction:** Despite the prevalence of shunt failure in the treatment of hydrocephalus and the near-constant need for shunt revisions, there are a limited number of methods that yield quick, quantifiable measures of catheter obstruction. We developed and assessed a gravity-driven device that measures flow through ventricular catheters. The model provides neurosurgeons with a quick, simple device that can send useful data to researchers. It can also be used by researchers to quantitatively compare catheter obstruction between different models of catheters. The model was used in this study to quantitatively analyze unused ventricular catheters used in the treatment of hydrocephalus; failed hydrocephalus catheters from our catheter biorepository were also quantitatively analyzed using the same device. The mechanisms of catheter obstruction are still poorly understood, but the literature suggests that resistance to fluid flow plays a significant role.

**Methods:** Catheters of three manufacturing companies were inserted into the benchtop model, which records time, flow rate, and pressure data using sensors. The relative resistances of catheters across six design models were evaluated. Experiments were performed to evaluate changes in the relative resistance of a catheter when the catheter’s holes were progressively closed. Relative resistance of explanted catheters from our catheter biorepository was also measured.

**Results:** Experimental results showed significant differences (*P* < 0.05) between the relative resistances of different catheter models just after being removed from their packaging. Furthermore, a trend of increasing resistance was observed in the experiments on catheters with manually plugged ventricular catheter holes. Data from five individual benchtop models were compared, and the differences in measured data between the models were found to be negligible. A significant increase (*P* < 0.05) in relative resistance was observed in explanted catheters.

**Conclusion:** The current study is meant both to validate the proposed model and to examine data on differences in relative resistance among catheter models. From these experiments, we can rapidly correlate clinical patient cohorts to identify mechanisms of luminal shunt obstruction. Collecting data for predictive analyses of potential patient outcomes is an area of potential future work, assuming sufficient sample size.

## Introduction

Hydrocephalus is a chronic disorder causing abnormal enlargement of the cerebral ventricles; the condition results from an imbalance in cerebrospinal fluid (CSF) production and absorption. The most common treatment for hydrocephalus is using surgically implanted shunts to drain excess CSF to other body parts, such as the peritoneal cavity. However, these shunts have notoriously high failure rates: approximately 50% of pediatric hydrocephalus shunts fail within two years of implantation.^1^ Virtually all hydrocephalus patients will require at least one shunt revision during their lifetime for various reasons ranging from infection to obstruction.^1,2^ Due to the high rate of catheter failure, often necessitating several surgical revisions over patients’ lifetimes, a great deal of scientific work has been dedicated to understanding the mechanisms of how and why catheters fail to use various approaches to improve catheters. The failure of catheters used to treat hydrocephalus is a multifaceted problem. Factors such as catheter hole configuration and the fluid dynamics of CSF flow through the catheters contribute to the problem.^3,4^

Much effort has been invested in modeling the fluid flow dynamics of catheters used to treat hydrocephalus. An equal amount of effort has investigated the effects of obstruction on CSF flow through these catheters. Two studies analyzed CSF flow through catheters; one of these studies examined several commercial catheter models. Both studies gave substantial evidence to suggest that the design of catheters plays a significant role in the obstruction and subsequent failure of hydrocephalus catheters.^3,4^ Another study used a hydrostatic pressure model to examine resistance in new and explanted catheters. This study found that there was some increase in catheter resistance over time. Explanted shunts also showed increases in resistance compared to unused catheters, but the literature was inconclusive on the clinical significance of these increases.^5^ There is limited work analyzing the effects of changes in resistance over time in catheters used to treat hydrocephalus and how these changes in resistance impact catheter obstruction. Further exploration of the relationships between implantation time, levels of catheter obstruction, and resistance to CSF flow would help supply context on the significance of the results reported in the literature.

Despite the incredible quantity of work on the subject, the mechanisms of shunt obstruction are still poorly understood. Further research on failed shunts is likely to help researchers continue expanding the knowledge base on shunt failure. To facilitate such research, a multicenter national biorepository was created at Wayne State University (WSU). From this biobank, a clinical database was created and used in a study to examine the effects of numerous factors on the number of revisions undergone by patients.^6^ A follow-up study imaged 343 of the WSU biobank’s ventricular catheters; these catheters were also classified depending on the number of holes occluded. Between the two studies, researchers characterized the tissue aggregates occluding the catheters; most failed catheters had obstructions.^7^ Such research has provided incredible insight into what factors affect catheter obstruction and catheter failure. An increased sample size would facilitate a better understanding of catheter obstruction and its symptoms. Thus, it could be useful to have a device that neurosurgeons and researchers can use to collect substantial amounts of quantitative data from catheters used to treat hydrocephalus.

The current study proposes a mass-producible, easy-to-use device that uses a water column and electronic sensors to acquire pressure, flow, and time data from ventricular hydrocephalus catheters. This device would be provided to neurosurgeons, who could then use the model to supply researchers with quantitative data from explanted shunts. These data could then be used to calculate and analyze resistance, allowing researchers to quantify and analyze changes in catheter performance over time. Two hypotheses were evaluated: (1) unused hydrocephalus catheters of different models will have significantly different resistances to fluid flow, and (2) resistance will increase as the catheter’s holes are closed. The model was validated by comparing data from individual 3D-printed devices without any catheters inserted to ensure differences in measured data were not significant.

## Materials and Methods

### The Design of the Benchtop Testing Device

The proposed design was designed for use in a hospital setting, and the priorities for its design were: (1) ease of use, (2) ability to quickly run successive samples, (3) mass producibility, and (4) portability. The proposed design utilized a water column to simulate a range of hydrostatic pressure similar to changes in intracranial pressure (ICP). The model, hereafter referred to as the Ventricular Catheter Testing Device (VCTD), was a simple in-and-out system with a position open for an in-line shunt catheter (Fig. 1). Water from the column flowed through the sample catheter before draining out through the silicone outlet tubing. CFD software was used to confirm that the flow through the catheter was laminar. Water flow was stopped and started using a plug that the operator screwed in or out. The model held a maximum of 10 mL of water. This volume of water was chosen to minimize the risk of damage to biobank samples while still providing a sufficient pressure drop for analysis. The CAD model (Fig. 1A) shows that the VCTD comprises three fused deposition modeling (FDM) 3D-printed components. The sample catheters were placed into a chamber made of UV-sensitive resin. A digital light processing (DLP) 3D printer was used to manufacture all components.

**Figure 1.**
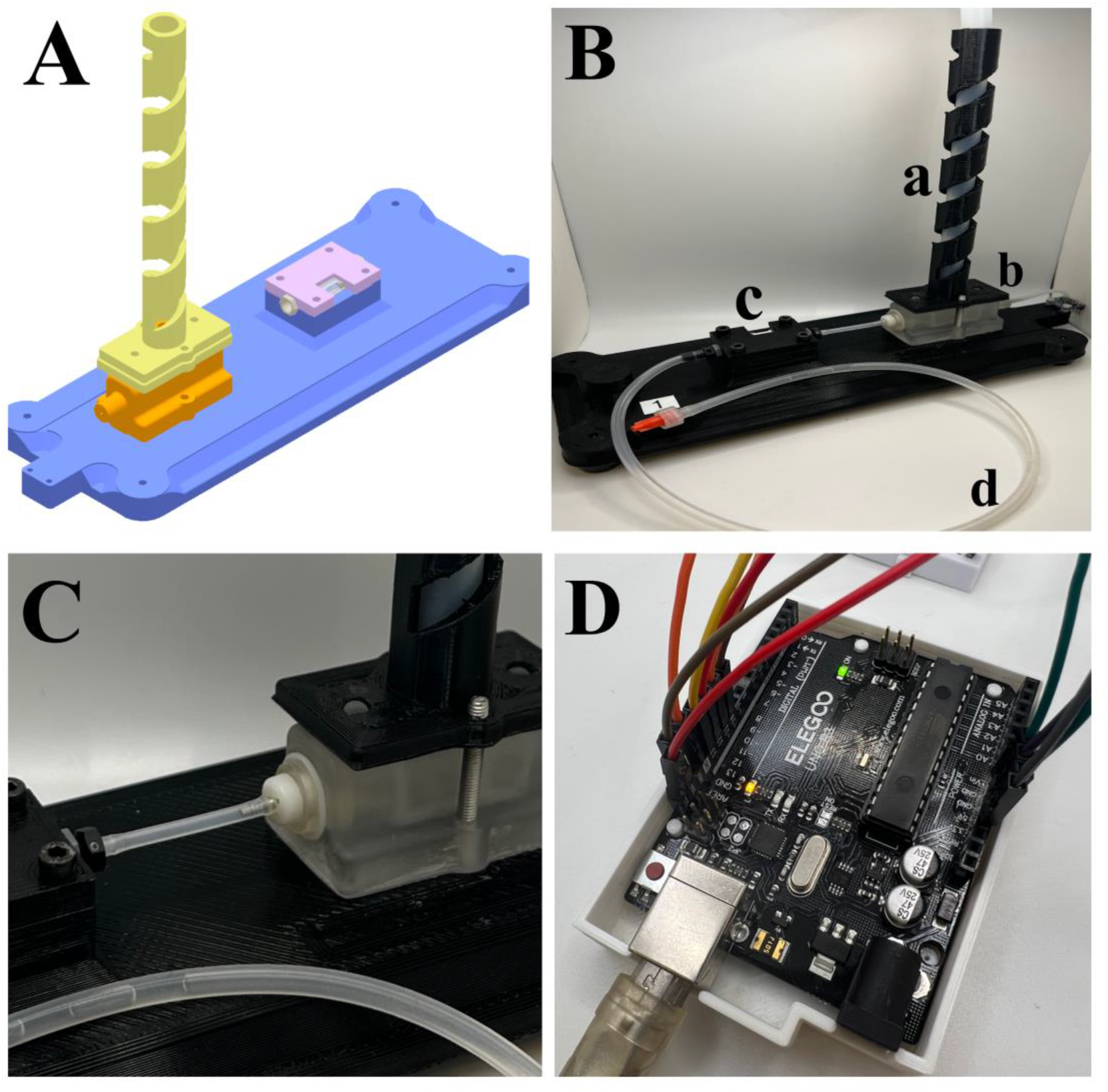
The Ventricular Catheter Testing Device. Figure 1A is a CAD model of the VCTD. The VCTD consists of the water column (yellow), the sample chamber (orange), and the sensor chamber (lavender), all attached to a single-piece base (blue). Figure 1B is an image of one 3D-printed VCTD. The major components of the VTCD are shown in Figure 1B: the water column (a), sample chamber (b), sensor chamber (c), and the silicone outlet tubing (d). Figure 1C is the sample chamber. After removing the silicone tubing, the sample holder can be removed from the chamber by turning the holder counterclockwise. Figure 1D is the AMD circuit board. The two lights visible are the yellow light and green light that notify the operator when the VCTD is ready to begin data acquisition. When the yellow light turns off and the green light stays on, the VCTD is ready to measure data from the sample. The VCTD shown on the right was used for all experiments testing catheters that were presented in this study.

A small circuit board component connected to the VCTD controlled data measurement and storage (Fig. 1D). The VCTD measures pressure in units of hPa and fluid flow in microliters/min. The circuit board used was an AMD ELEGOO UNO R3 Arduino-compatible board manufactured in California. The VCTD was designed for automated data acquisition: tasks for the operator included inserting and removing the sample catheter, opening and closing the flow valve, and filling the water column. Data were automatically obtained from pressure sensors (Adafruit MPRLS Ported Pressure Sensor, 0-25 PSI, manufactured in New York) and flow sensors (Sensirion AG SLF3S-1300F, 40 mL/min, manufactured in Switzerland) positioned directly after the catheter. The pressure of the water column was similarly measured using a pressure sensor at the bottom of the column. The pressure sensor also measured the atmospheric pressure to account for differences in testing site altitude. All sensors sent data to a SanDisk Ultra 16GB Micro SDHC Class 10 chip, which stored the data as a comma-separated values (csv) file. These files were then imported into Microsoft Excel.

### Procedure for Testing Catheters Using the VCTD

The procedure for using the VCTD was straightforward and was as follows: first, the sample catheter was cut to a length of 40 millimeters using the groove on the side of the VCTD. Next, the tubing that connects the flow sensor to the sample holder was removed. The sample holder (Fig. 1C) was unscrewed by turning it counterclockwise until it detached from the device. The sample catheter was eased onto the sample holder, ensuring that the sample was securely and entirely on the sample holder. Next, the sample holder was screwed back into the VCTD by turning the holder clockwise, and the previously removed tubing was reattached. After this, the flow was shut off using the clamp to pin down the end of the tubing. The column was filled with water up to the top using a syringe. The column was carefully filled to avoid trapping air bubbles at the bottom of the column. If any trapped air bubbles were visible, the top of the column was covered, and the bottom was firmly tapped until bubbles were dislodged. Once the column was filled, the device was plugged into an outlet. Before continuing, the operator waited for the green light to turn on and for the yellow light to turn off (Fig. 1D).

Once the yellow light turned off, the device was ready to collect data; the plug was removed to allow fluid to flow. The VCTD was not disturbed while the water was flowing. The operator waited five seconds after flow ceased before unplugging the VCTD to stop data acquisition. The tubing was removed from the sample holder, the sample holder was unscrewed, and the sample catheter was removed. After removal of the catheter, the sample holder was wiped using a Kimwipe and 60% w/v ethanol. Once the ethanol had evaporated fully, another sample would replace it. If there were no other samples for testing, the empty sample holder was screwed back into the VCTD, and the tubing was reattached. No particular containment procedure was needed to store the VCTD, but it was ensured that the electronic components avoided contact with fluids. After use, the VCTD was allowed to air dry. Distilled, deionized water was used for all experiments and for rinsing the VCTD’s tubing.

### Experimental Hypotheses and Methods

We performed four categories of experiments: (1) experiments testing five individual VCTD devices with no catheters inserted, (2) experiments with unused catheters inserted into the VCTD, (3) experiments with patient-explanted catheters inserted into the VCTD, and (4) experiments with an unused catheter that had its holes progressively closed inserted into the VCTD. All experiments with catheters were performed using VCTD #1 (n = 5 for all experiments).

We created a live catalog of commercial catheters to help select catheters for testing. The catalog listed catheter and valve products used to treat hydrocephalus from several American biomedical manufacturing companies. Products were categorized by name, reference number, and manufacturer. Product details such as device length, internal diameter, outer diameter, and hole configuration were included, if available. Unused, unexpired catheters were acquired directly from manufacturers (via eSutures). Two different models from three different biomedical manufacturing companies were tested, totaling six different ventricular hydrocephalus catheter models. We selected the models due to availability, dissimilarity, and brand name recognition. The six tested models were labeled in this study by generic labels: Manufacturer 1 Model 1 (M1M1), Manufacturer 2 Model 1 (M2M1), and so on. Explanted hydrocephalus catheters were obtained from Wayne State University’s biobank of consented neurosurgical samples. Explanted samples were labeled as Explanted 1, Explanted 2, *et cetera*. Five explanted samples across two centers were tested. Samples were moved using sterile tweezers; gloves were worn during handling at all times. The measuring surface and the tweezers were cleaned with 60% w/v ethanol after each sample was moved.

The experiments measuring flow through the VCTD without any inserted catheter were performed on five different VCTD models (n=5). The unused catheter with plugged holes was obstructed manually in the lab using transparent, Gorilla-brand hot glue sticks manufactured in Ohio. A new M1M1 catheter was obstructed by using a hot glue gun to apply a thin layer of glue to each row of holes. Care was taken to ensure the hot glue gun’s tip did not contact the catheter. The catheter was not heated appreciably during the application of glue. Firm pressure was applied to the glue-coated surface while the glue was cooling to ensure the plugging of the holes. Once the glue had fully solidified, the catheter was placed inside the VCTD for testing. Data for the manually obstructed catheter were categorized based on how many rows of holes were plugged with glue. For example, data for the unobstructed catheter were labeled as 0RO (i.e., Zero Rows Obstructed).

### Data Curation and Analysis

Once pressure, flow, and time data were acquired, the hydrostatic pressure resulting from the water column at each data point was calculated by subtracting the end pressure (i.e., atmospheric pressure) from the total pressure at each data point. A range of pressures, from 15 hPa to 7 hPa, was selected to ignore noise observed at the beginning and end of each experiment (Fig. 2). The chosen range used approximately 600 data points.

**Figure 2.**
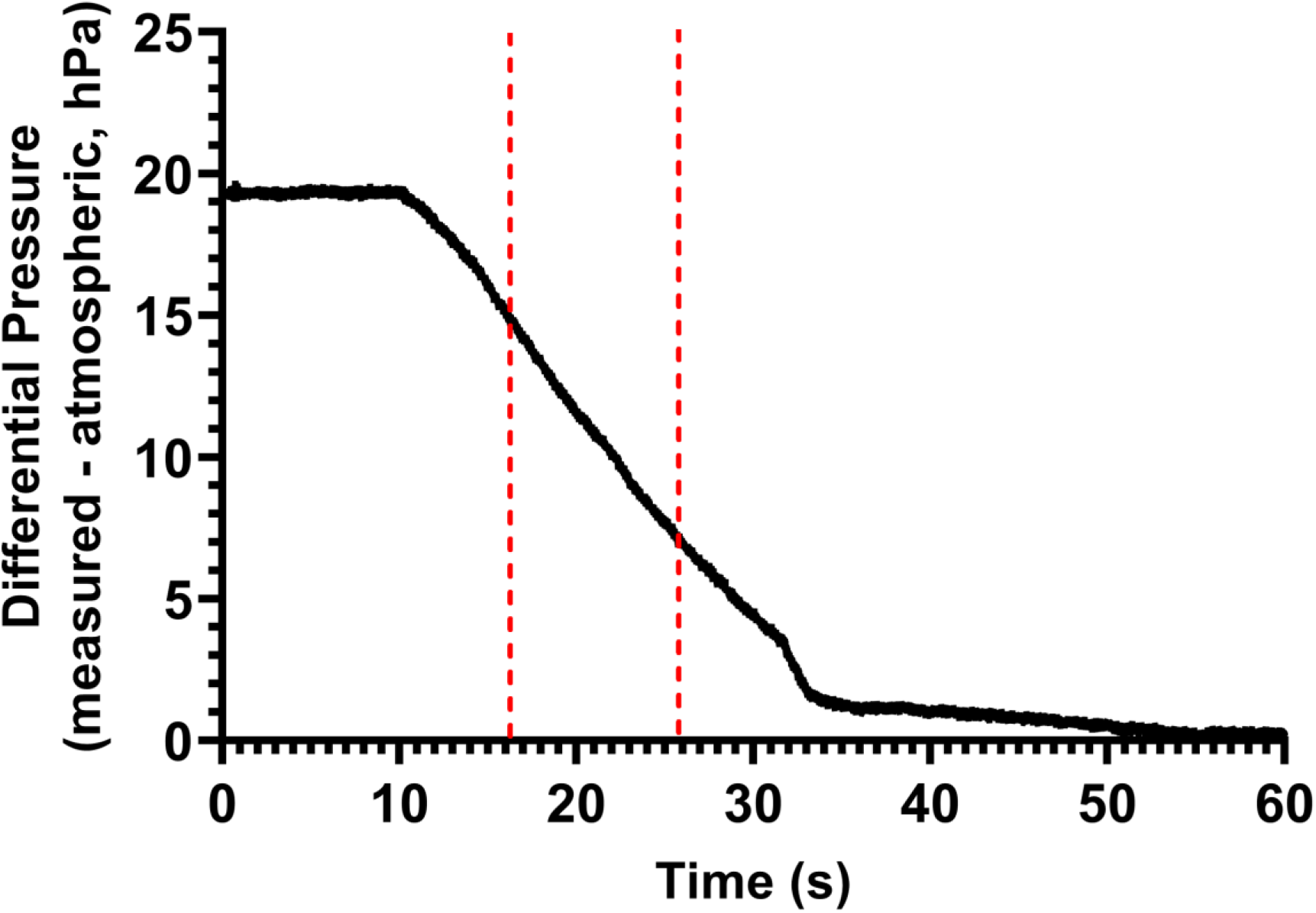
Representative pressure vs time graph for an unused M1M1 catheter. Sample of data acquired using the VCTD. Data is from one experimental run of an unused hydrocephalus catheter of model M1M1. The Y-axis shows the hydrostatic pressure, which was calculated by subtracting atmospheric pressure from total pressure at all points. The section of data enclosed by the red, vertical lines was used for data analysis. The enclosed data region represents the time elapsed for hydrostatic pressure to drop from 15 hPa to 7 hPa. The time elapsed from 15 hPa to 7 hPa was used for all analyses.

Once the hydrostatic pressure was found for the dataset, the time elapsed from 15 hPa to 7 hPa was calculated. This calculation was performed for each run, and then an average time elapsed (in seconds), and the standard deviation was calculated for each experimental group. These average time elapsed and standard deviation data were used for statistical analysis.

Time elapsed was used to compare differences in resistances between groups. The relationship between pressure, flow rate, and resistance for a Newtonian fluid through a tube is well known (Eq. 1).^8,9^ The pressure drop for each experiment was approximately the same; hence, differences in flow rate between catheters resulted from differences in resistance. These resistances, inferred from the time elapsed for a fixed pressure drop, were termed “relative resistances.” A longer time elapsed meant a greater resistance to flow, and a shorter time meant a lesser resistance to flow.

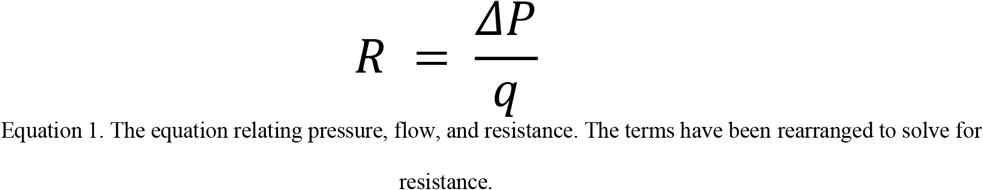

Pressure and flow data were used to calculate resistance values to validate the use of relative resistance as a metric for comparing resistances between catheters. An excellent study by Cheatle et al. calculated resistances for catheters implanted in hydrocephalus patients.^5^ Resistance values were calculated using data from the VCTD and compared to the resistance values reported in literature. Pressure vs. flow graphs were constructed for each dataset (Fig. 3) to calculate resistances with VCTD data. Linear regression was performed on each dataset using these graphs. The relationship between pressure, flow, and resistance (Eq. 1) was used to determine that the slope of a given sample’s pressure vs. flow graph was the resistance for that sample. The slope and R^2^ values for each linear regression were calculated and recorded. A dimensional analysis was performed to convert from units of hPa•min/µL to Pa•s/m^3^ to match units with the literature.

**Figure 3.**
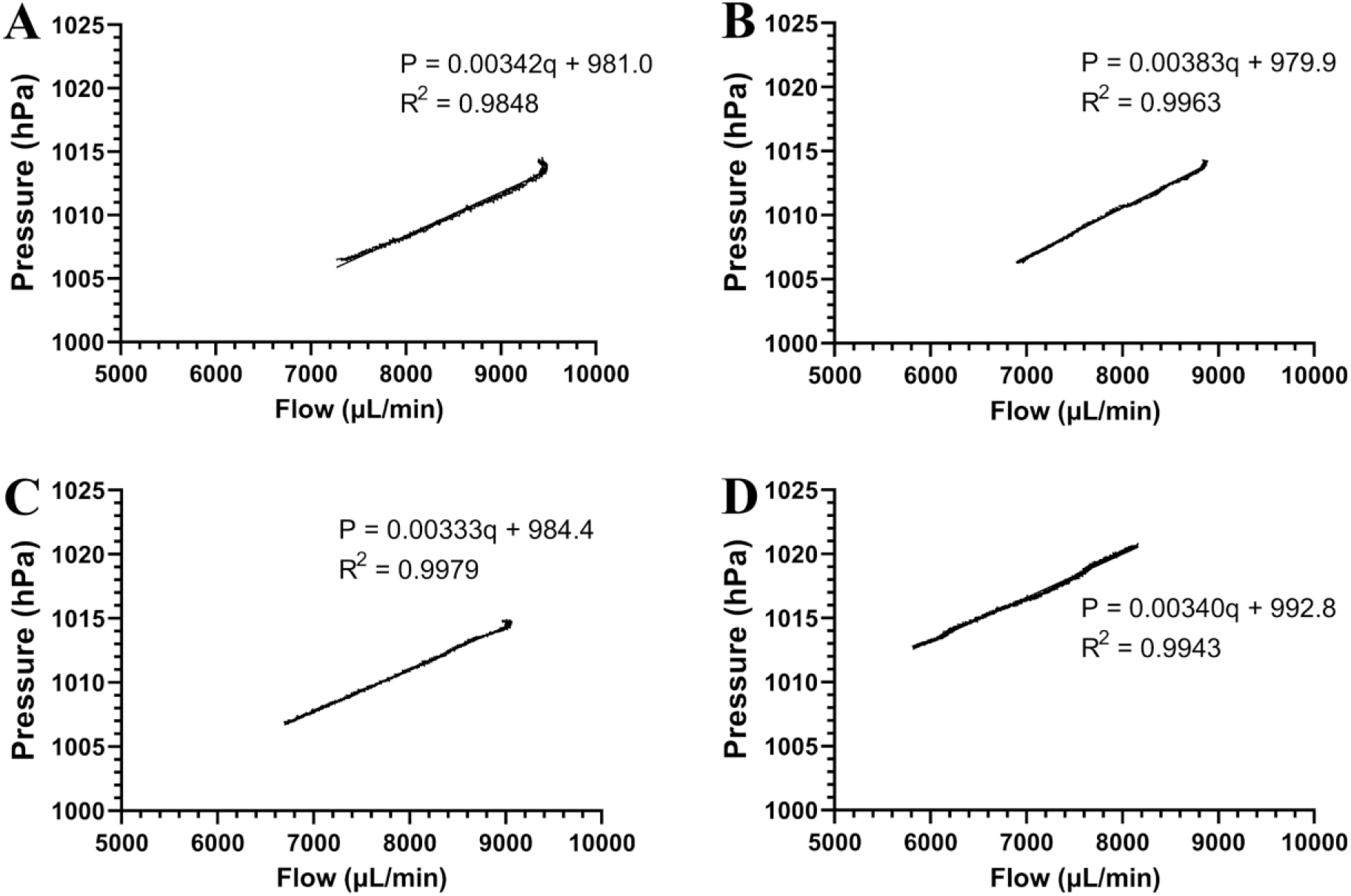
Examples of pressure vs flow graphs created using raw data taken by VCTD #1. The graphs are for the VCTD alone (3A), the VCTD with an unused M1M1 catheter inserted (3B), the VCTD with an explanted catheter inserted (3C), and the VCTD with the 4RO manually obstructed catheter inserted (3D). A rolling average of 110 data points was used for the flow data when constructing these graphs; the rolling average was used to reduce noise from data measurement. The slope of the linear regression equations for each graph was used to calculate resistance.

### Statistical Analysis

Microsoft Excel for Windows was used to process data (e.g., calculating each time elapsed and calculating hydrostatic pressure). Statistical analyses were performed using SPSS v28 for Windows and an alpha value of 0.05. Time elapsed data were analyzed for normality by constructing histograms and visually comparing data to a normal distribution curve. A one-way analysis of variances (ANOVA) test was used to determine significant differences between data of a given experimental group; a Tukey post hoc test followed each ANOVA. Box-and-whisker plots were constructed using SPSS; all other graphs were made using GraphPad Prism version 9 for Windows. Separate hypothesis categories were not compared to each other. For example, data from the unused catheter with progressively plugged holes were not compared to data from the other unused catheters.

## Results

### Manually Obstructed Catheter

The results for the manually obstructed catheter (Table 1) showed that the differences in time elapsed were significant (*P* < 0.001). Post-hoc analysis (Fig. 4A) further showed that the time elapsed for the four-rows-plugged catheter was significantly different from the time elapsed for all other groups (*P* < 0.001). Furthermore, the time elapsed for the unplugged catheter and one-row-plugged catheter were significantly different compared to the time elapsed for the three-rows-plugged catheter (*P* < 0.05). The time elapsed data for the unplugged catheter, one-row-plugged catheter, and two-rows-plugged were not significantly different (*P* > 0.05). Similarly, the time elapsed data for the two-rows-plugged and three-rows-plugged catheters were not significantly different (*P* > 0.05). A trend of increasing relative resistance was observed as holes were plugged; the trend appears to be exponential (Fig. 4B).

**Table 1.**
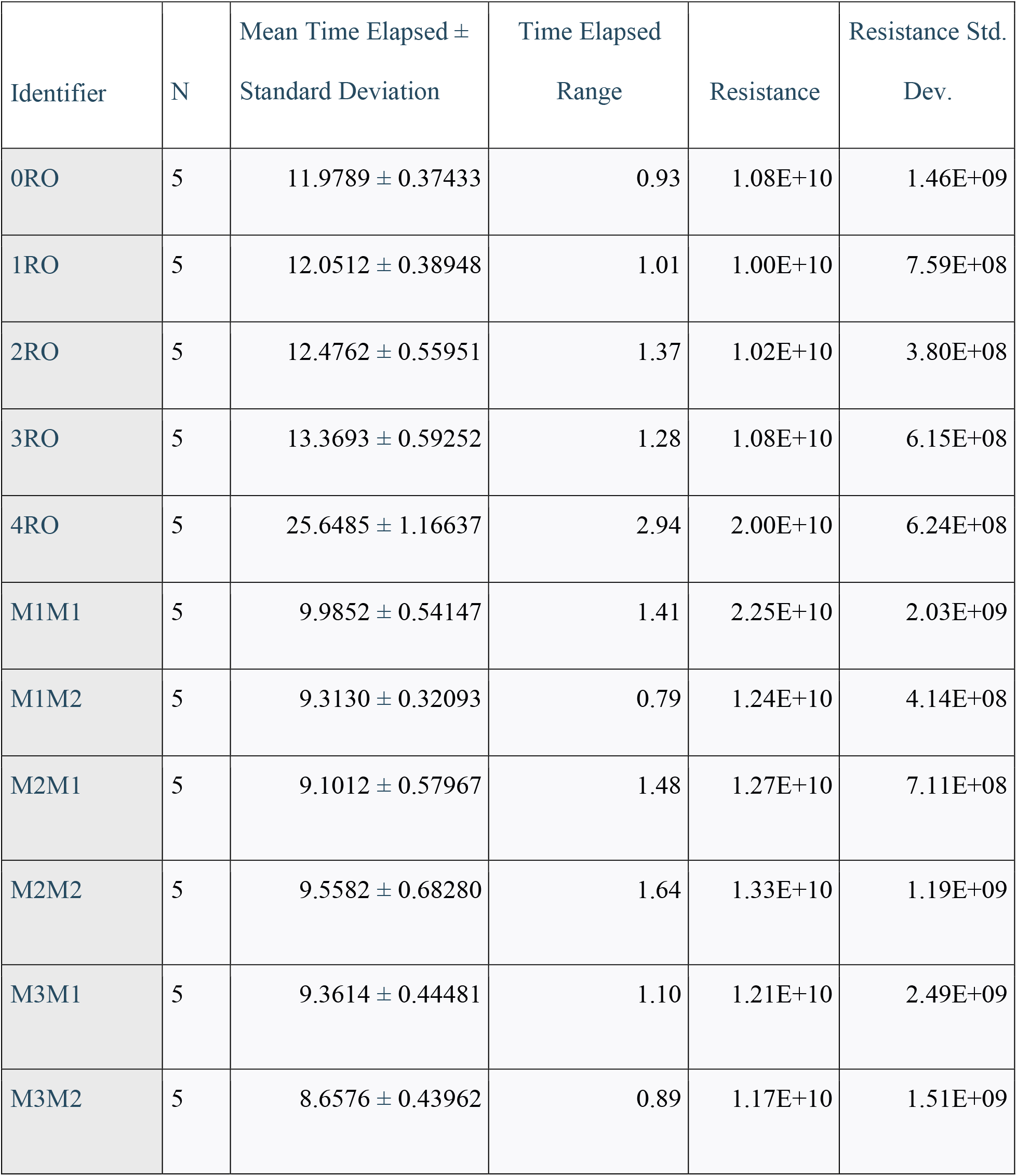
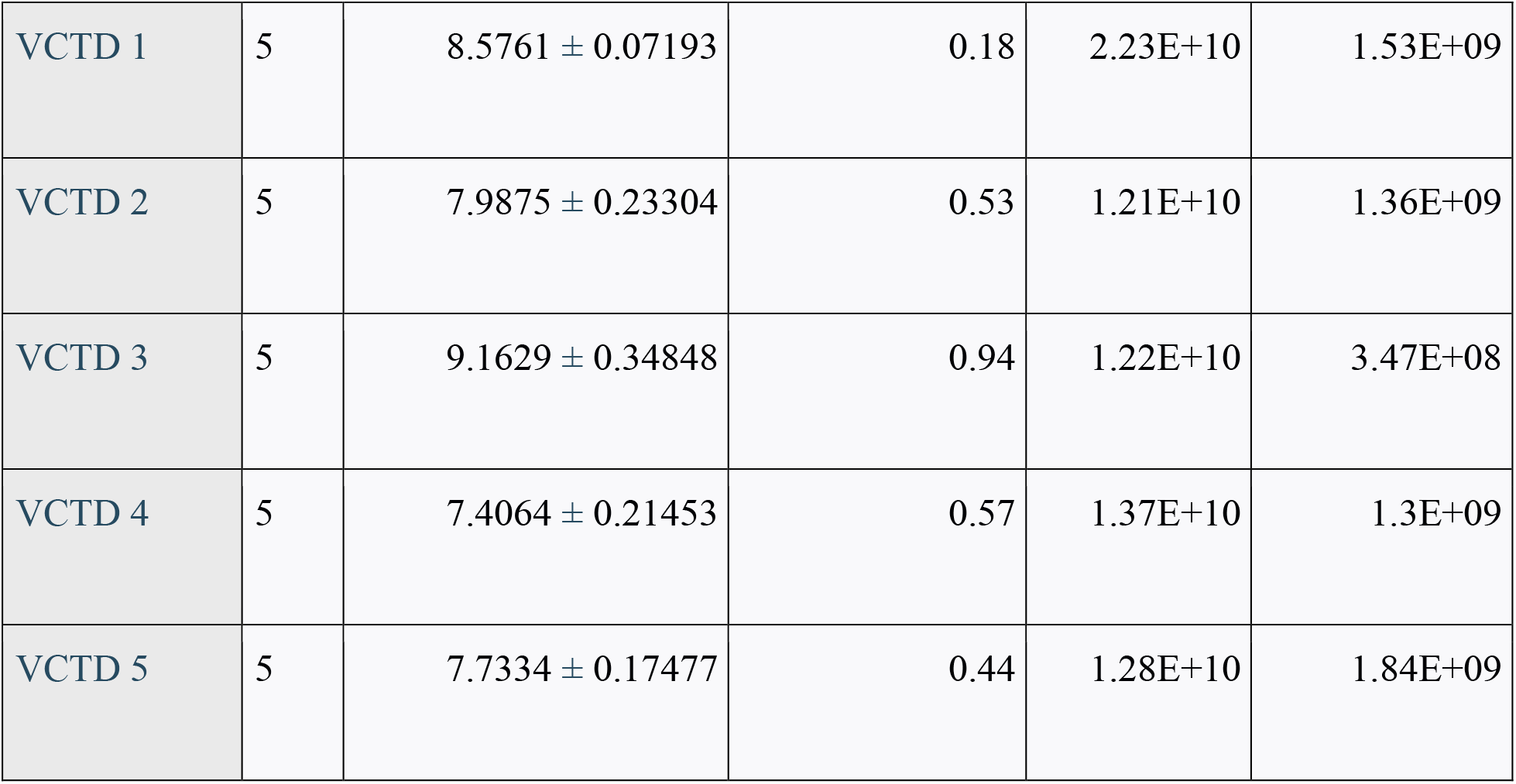
Table displaying n, mean time elapsed with standard deviation, time elapsed range, average calculated resistance, and resistance standard deviation for all experimental groups presented in this study. Mean time elapsed and time elapsed range have units of seconds. Resistance and resistance standard deviation have units of Pa•s/m^3^.

| Identifier | N | Mean Time Elapsed ±<br>Standard Deviation | Time Elapsed<br>Range | Resistance | Resistance Std.<br>Dev. |
| --- | --- | --- | --- | --- | --- |
| 0RO | 5 | 11.9789 ± 0.37433 | 0.93 | 1.08E+10 | 1.46E+09 |
| 1RO | 5 | 12.0512 ± 0.38948 | 1.01 | 1.00E+10 | 7.59E+08 |
| 2RO | 5 | 12.4762 ± 0.55951 | 1.37 | 1.02E+10 | 3.80E+08 |
| 3RO | 5 | 13.3693 ± 0.59252 | 1.28 | 1.08E+10 | 6.15E+08 |
| 4RO | 5 | 25.6485 ± 1.16637 | 2.94 | 2.00E+10 | 6.24E+08 |
| M1M1 | 5 | 9.9852 ± 0.54147 | 1.41 | 2.25E+10 | 2.03E+09 |
| M1M2 | 5 | 9.3130 ± 0.32093 | 0.79 | 1.24E+10 | 4.14E+08 |
| M2M1 | 5 | 9.1012 ± 0.57967 | 1.48 | 1.27E+10 | 7.11E+08 |
| M2M2 | 5 | 9.5582 ± 0.68280 | 1.64 | 1.33E+10 | 1.19E+09 |
| M3M1 | 5 | 9.3614 ± 0.44481 | 1.10 | 1.21E+10 | 2.49E+09 |
| M3M2 | 5 | 8.6576 ± 0.43962 | 0.89 | 1.17E+10 | 1.51E+09 |
| VCTD 1 | 5 | $8.5761 \pm 0.07193$ | 0.18 | 2.23E+10 | 1.53E+09 |
| VCTD 2 | 5 | $7.9875 \pm 0.23304$ | 0.53 | 1.21E+10 | 1.36E+09 |
| VCTD 3 | 5 | $9.1629 \pm 0.34848$ | 0.94 | 1.22E+10 | 3.47E+08 |
| VCTD 4 | 5 | $7.4064 \pm 0.21453$ | 0.57 | 1.37E+10 | 1.3E+09 |
| VCTD 5 | 5 | $7.7334 \pm 0.17477$ | 0.44 | 1.28E+10 | 1.84E+09 |

**Figure 4.**
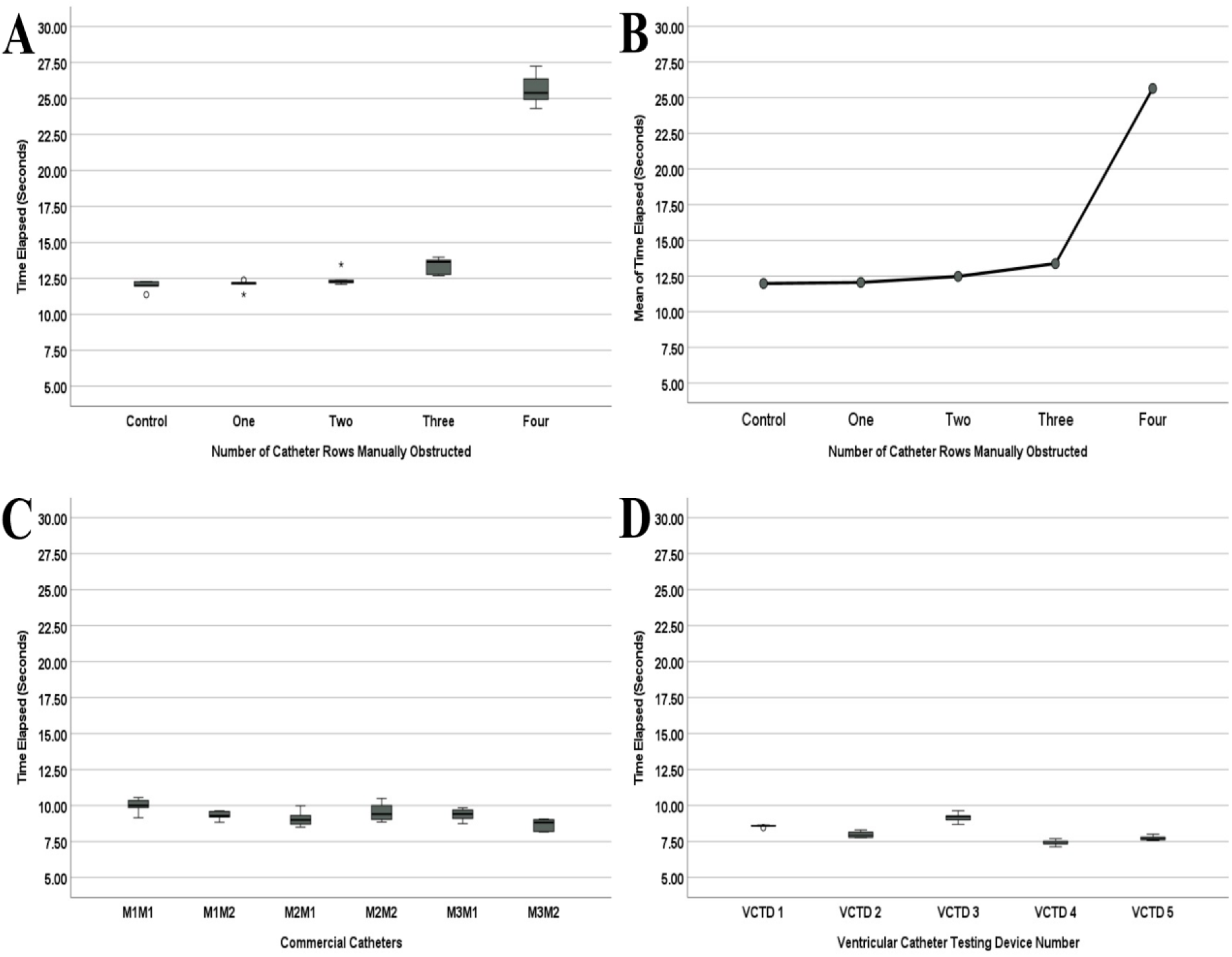
Collage of graphed experimental results. Figure 4A is the box-and-whiskers plot of the mean time elapsed data for the manually obstructed catheter. Figure 4B shows the trend in relative resistance between the means of the groups for the manually obstructed catheter experiments. Figure 4C is the box-and-whiskers plot of the mean time elapsed data for the unused commercial catheters. Figure 4D is the box-and-whiskers plot of the mean time elapsed data for the individual VCTDs. All mean time elapsed data is reported in seconds (n=5).

### Commercial Catheter Comparison

The results of the experiments on commercial catheters (Table 1) showed that there were significant differences between the time elapsed data for different models (*P* < 0.05). Post-hoc analysis (Fig. 4C) showed that the differences between time elapsed data for only two catheter models, M1M1 and M3M2, were significantly different (*P* = 0.005). Conversely, the time elapsed data between all other model pairs were insignificant (*P* > 0.05).

### Comparison of Data Measurement Among Individual VCTDs

Each VCTD was tested five times using the procedure described in the Methods section. No catheters were inserted into any of the VCTDs during these experiments. The results (Table 1) found that the differences between the time elapsed data for each VCTD were significant (*P* < 0.001). The post-hoc analysis (Fig. 4D) showed that VCTD #1 was significantly different from all other Set Ups (*P* < 0.005). Furthermore, VCTD #2 was significantly different from VCTD #3 (*P* < 0.001) and VCTD #4 (*P* = 0.005). Finally. VCTD #3 was significantly different than VCTD #4 (*P* < 0.001). VCTD #2 and VCTD #5 were not significantly different (*P* > 0.05). The data for VCTD #4 and VCTD #5 were also not significantly different (*P* > 0.05).

### Calculation of Catheter Resistance using Pressure and Flow Data

All of the linear regressions performed on the data presented in this study had an R2 value between 0.99 and 1.00. The calculated resistance values (Table 1) were compared to the relative resistances. It was found that a greater calculated resistance value generally correlated with a greater relative resistance (i.e., a higher time elapsed). The only exceptions to this observed trend among the catheters presented in this study were 0RO and M2M1; despite having lower relative resistances compared to other samples in their respective groups, these two samples had higher calculated resistances compared to the other samples in those groups. VCTD #1 also shows higher calculated resistance than VCTD #3, despite having a lower time elapsed.

## Discussion

Significant differences were found between the individual VCTD units’ relative resistances. However, all the experiments with catheters presented in this study were performed using the same VCTD; variations between VCTDs will not have impacted data acquisition. Furthermore, the standard deviation and range for each VCTD are small (Table 1). Some degree of variance between models is inevitable, even with 3D printing. The base relative resistance of every VCTD will be measured and recorded to account for this variance. As such, it is unlikely that the variations in data measurement observed in this study will cause significant problems for data analysis in future studies that use VCTDs for data acquisition.

### The Use of Relative Resistance to Infer Resistance to CSF Flow

The values of resistance calculated in this study are similar to those reported in literature.^5^ Cheatle et al. suggested that increases in the resistance of catheters could contribute to shunt failure by reducing the flow rate of CSF through the shunt system.^5^ While the catheters presented in this study cannot be directly compared to those presented by Cheatle et al., there is strong evidence to suggest that relative resistance is connected to the catheters’ resistance. Thus, we might suspect that the VCTD can compare changes in resistance to CSF flow using relative resistance. Indeed, the data presented in this study suggest that catheters with a higher relative resistance have a lesser flow rate, evidenced by the higher time interval required for a fixed pressure drop. Given that the results of this study suggest that relative resistance can be used to infer catheter resistance, we suspect that changes in relative resistance could be used to quantitatively categorize decreases in catheter performance caused by increased resistance to CSF flow. Factors such as tissue obstruction and catheter geometry can influence CSF flow through catheters. The results of this study present evidence to link changes in relative resistance to such factors.

### Preliminary Analysis of Explanted Catheters from Patients with Hydrocephalus

Utilizing biobanks to analyze a sufficiently large quantity of samples can help account for both the heterogeneity between patients and the broad range of hydrocephalus etiologies.^6,7^ Integrating the VCTD when analyzing biobank samples may provide researchers with valuable data that could help characterize shunt obstruction, examine factors that affect shunt failure, and analyze the efficacy of commercial catheters. An analysis of the samples at the WSU biorepository found that a majority (61.5%) of the failed shunts contained obstructive tissue aggregates.^7^ Several explanted catheters presented in this study show significant differences in relative resistances (Fig. 5).

**Figure 5.**
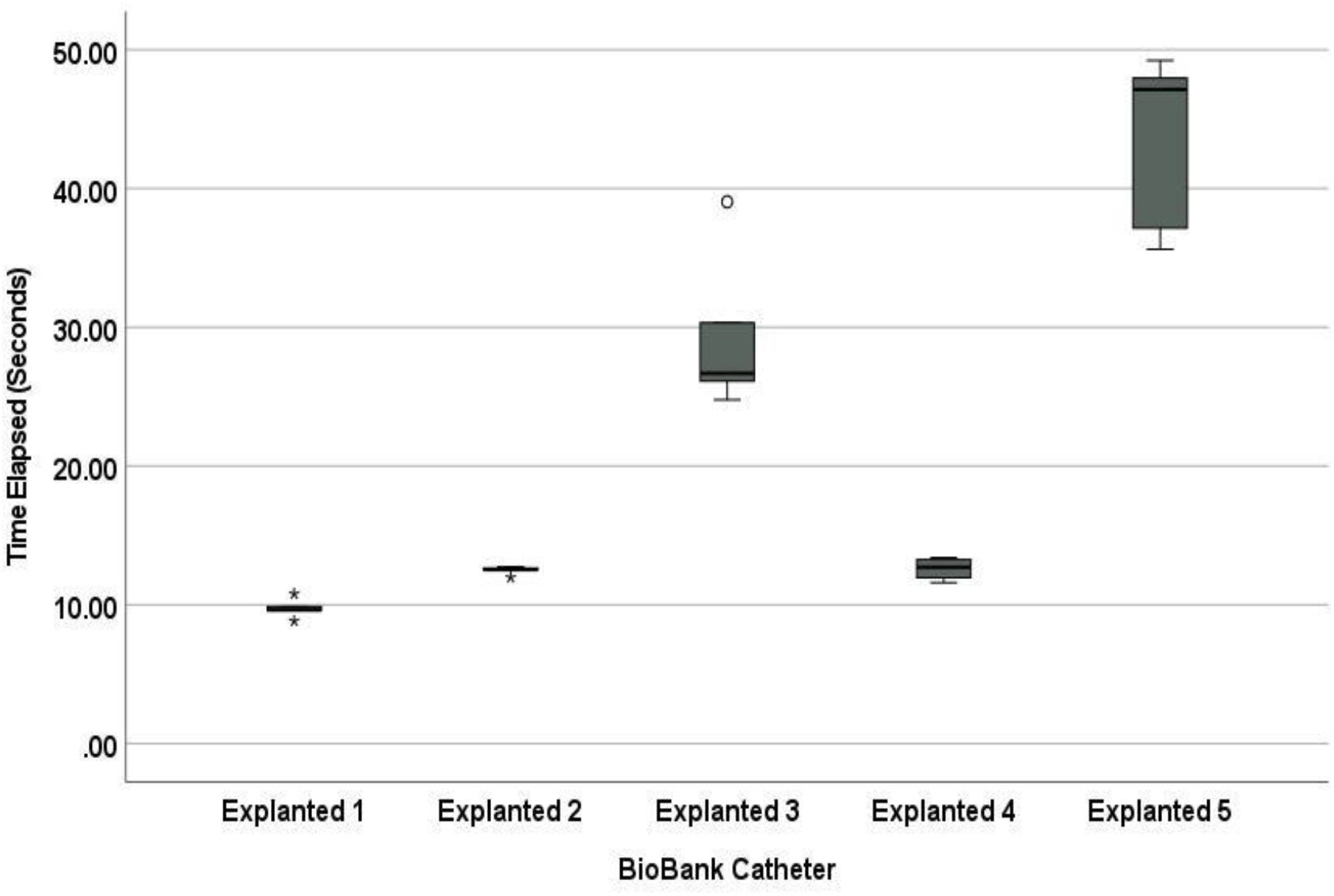
Box-and-whiskers plot of the time elapsed data for the explanted biobank catheters (n=5). Asterisks and circles represent data points that are outliers.

The prevalence of tissue aggregates and the noticeable increase in relative resistance compared to unused catheters (Table 1, Table 2) strongly suggest a significant relationship between obstruction by tissue aggregates and increased resistance to CSF flow. However, the physiological significance of the biobank data is still unclear. A significant limitation in analyzing these biobank samples is that the models of many of the explanted catheters are uncertain. Thus, these explanted samples cannot be directly compared to individual unused catheters. The catalog of catheters seeks to address this issue; physical characteristics such as hole size and configuration could be used to determine the model of biobank shunts. Identifying the models of the biobank samples would facilitate direct comparisons of the relative resistances of different catheter models before and after implantation. Furthermore, a couple of the tested explanted catheters are close in relative resistance (Table 2); a simple visual examination of the hole configurations for these catheters suggested they might be of the same model. The ability to confirm the models of the biobank samples would thus also enable comparisons of relative resistance between failed and unused catheters of the same model.

**Table 2.**
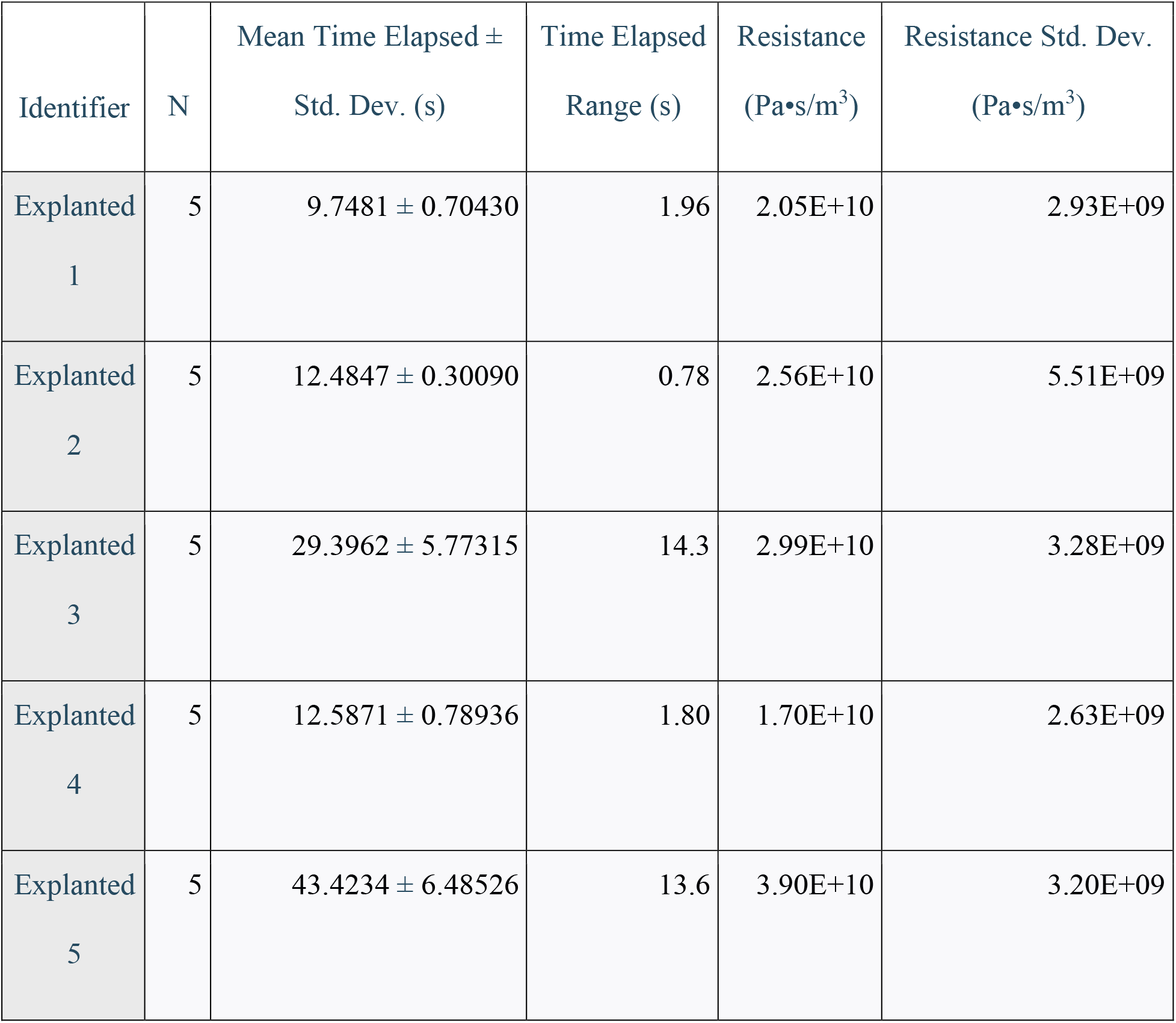
Results for biobank catheters. Mean time elapsed and time elapsed range have units of seconds. Resistance and resistance standard deviation have units of Pa•s/m^3^.

Once such comparisons can be performed, researchers can establish metrics of resistances for catheters, ranging from unused to high risk of failure. Various factors have been used to predict shunt failures, such as shunt placement, patient age, CSF composition, and the number of neurosurgeons operating on the patient.^10–13^ Data on changes in catheter resistance over time could provide another parameter for these predictive analyses. Expanding on the shunt catalog would also provide a convenient method for researchers to share data on various catheter designs. The creation and upkeep of the shunt catalog is a long-term project involving the VCTD and will be used for several future studies.

### Relationships Between Catheter Geometry, Hole Obstruction, and Resistance

The post-hoc analysis of data from the manually obstructed catheter showed that the most significant difference in relative resistance occurred when all four rows were manually obstructed. There were no significant differences in relative resistances compared to the unobstructed catheter until three rows of holes (i.e., more than half) were plugged. Hence, there is evidence to suggest that the effect of obstruction on resistance to fluid flow is nonlinear. Not all catheter models have four rows of holes with eight holes per row; future testing will examine if this apparent nonlinear relationship holds for different hole configurations. If further evidence is found to support this nonlinear relationship, it would suggest that signs of obstruction experienced by patients occur after the implanted catheter is almost entirely obstructed. Further investigations on progressive catheter obstruction could help better characterize this relationship to predict when shunt obstruction will contribute to failure.

There have been several investigations on the mechanisms of catheter obstruction. Increased flow and shear stress have been observed to promote cell adhesion counterintuitively.^14,15^ CSF flow through the lumen of catheters induces shear stress on astrocytes and macrophages, causing them to adhere to the inner walls of the catheters. This phenomenon is known to contribute to the catheter lumen’s obstruction significantly.^15–17^ However, literature has also observed tissue growths forming on the catheter’s surface, infiltrating the catheter holes, and then infiltrating the lumen.^7,17–18^ CSF flows through the catheter’s holes before entering the lumen, thus making the holes susceptible to cell adhesion and obstruction. Studies have manipulated catheter hole size, hole number, and hole placement to examine the effects on cell adhesion.^3,16–17,19^ The results of these experiments suggest that changing the geometry of the catheters themselves may substantially help reduce shunt obstruction. Despite the incredible, extensive work on the subject, there is still much to learn about the mechanisms underlying catheter obstruction. Further investigation of the mechanisms by which shear stress affects cell adhesion, the differences between catheter obstruction originating in the lumen versus obstruction originating from the outside of the catheter, and the relationship between obstruction and resistance to CSF flow is necessary. Future work using the VCTD in a controlled study, examining differences in relative resistance between obstructed catheters and unobstructed catheters with various hole configurations, could facilitate these investigations into the factors affecting catheter resistance and the subsequent effect of resistance on catheter obstruction. These studies would be very beneficial in understanding the clinical significance of changes in catheter resistance.

### Differences in Relative Resistance Among Commercial Models

The data acquired in this study indicate a significant difference in relative resistances among catheter manufacturers and models. However, the clinical significance of these differences is unclear. There is limited evidence in literature to suggest any relationship between clinical outcomes and the model of catheter used. The relative resistances between most commercial catheters tested in this study were not significantly different. The study on commercial catheters performed by Galarza et al. found that flow distribution followed similar patterns across the commercial catheters they tested. Their results suggested that hydrocephalus catheters might fail with some degree of uniformity.^4^ From a fluid dynamics standpoint, catheters of the same model had similar flow characteristics and experienced obstruction similarly.^4^ Future work using the VCTD will determine whether there is significant variation between the same model’s relative resistances of individual catheters.

### Potential Improvements in the VCTD’s Design

A significant limitation of the VCTD is that it cannot represent the unique conditions that each shunt experiences inside a patient. As shown in the biobank studies by Gluski et al. and Hariharan et al., many factors are related to shunt revision.^6,7^ Each patient’s unique biological environment and personal lifestyle will also affect shunt performance. Measurements taken by the VCTD also do not account for differences in the physical dimensions of catheter models. Commercial catheter models boast various catheter diameters, hole configurations, and lengths. Previous research has established clear relationships between these parameters and the flow of CSF through a catheter.^3–5,14–17,19^ There is also the fact that the removal of shunts from a patient is likely to affect tissue aggregates present, such that the sample tested is not in the same condition as it was prior to removal. Many of the samples in the WSU biobank underwent electrocautery during the removal process, which may have affected the tissue or the catheter itself. The biobank samples tested were also fixed for imaging, which would also have impacted the tissue. Concerning the latter limitation of the biobank samples, one benefit of the VCTD is that fixing is unnecessary to run experiments. Explanted shunts can be tested immediately after removal. However, the other limitations are unavoidable and encountered by most *in vitro* research that models hydrocephalus. Future work using the VCTD will need to account for the limitations present in this study.

Data acquired from the VCTD cannot currently be used to predict whether a shunt at a given resistance will fail. However, the data can be used to characterize and compare levels of obstruction. The data from the VCTD can be used to calculate resistance, but more samples need to be tested to validate the resistance calculations fully. The VCTD is not a bioreactor; it does not simulate flow patterns in the brain. The VCTD is a gravity-driven model, which is not equivalent to how CSF flows through the body. Benchtop pump systems can simulate CSF flow with varying parameters and high precision and have been proposed as potential components of *in vitro* hydrocephalus models.^20^ Finally, the manually obstructed catheter was obstructed using glue, which is notably different from being obstructed by human cells. Future experiments with manually obstructed shunts can use human astrocyte cultures to plug the hole interfaces, providing a more accurate shunt obstruction model. These approaches could help researchers better model the relationship between resistance to CSF flow and catheter obstruction. Integrating these approaches for future iterations of the VCTD would address some limitations.

## Conclusions

This study used a hydrostatic benchtop model to simulate fluid flow through unused and explanted catheters used to treat hydrocephalus. Experiments were also performed on a catheter that had been manually obstructed and on the benchtop model itself. The results of the experiments showed significant differences in relative resistance between some commercial models and significant differences in relative resistance when an unused catheter’s hole interfaces are closed. A non-linear increasing trend in relative resistance was observed as the hole interfaces were progressively closed. Differences in data between individual benchtop models were found to be significantly different. However, these differences are unlikely to affect data measurement and analysis meaningfully due to the small standard deviation within each model’s measurements and the fact that each model will be calibrated prior to use.

Literature shows that the fluid flow characteristics and geometry of catheters contribute to shunt obstruction, which is responsible for most shunt failures. The mechanisms related to shunt obstruction and failure are still poorly understood. The VCTD seeks to provide researchers and neurosurgeons with a device that can facilitate the rapid collection and analysis of substantial amounts of clinical data with high resolution. This study and the proposed VCTD model are meant to be foundations for fulfilling several long-term goals: (1) the quantitative characterization of bulk catheter obstruction and inhibition of CSF outflow through the catheter, (2) the establishment of a catalog of commercial catheters and their subsequent long-term performance, (3) a method by which surgeons can quickly and easily provide large amounts of quantitative data from explanted shunts, (4) an analysis of the relationships between resistance and ICP as an indirect function of CSF production, and (5) analyses on the efficacies of commercially available catheters used to treat hydrocephalus.

## List of Abbreviations

CSF: Cerebrospinal Fluid
CNS: Central Nervous System
CFD: Computational Flow Dynamics
WSU: Wayne State University
ICP: Intracranial Pressure
FDM: Fused Deposition Modeling
DLP: Digital Light Processing
VCTD: Ventricular Catheter Testing Device
M1M1: Manufacturer 1 Model 1
M1M2: Manufacturer 1 Model 2
M2M1: Manufacturer 2 Model 1
M2M2: Manufacturer 2 Model 2
M3M1: Manufacturer 3 Model 1
M3M2: Manufacturer 3 Model 2
0RO: Zero Rows Obstructed
1RO: One Row Obstructed
2RO: Two Rows Obstructed
3RO: Three Rows Obstructed
4RO: Four Rows Obstructed
ANOVA: Analysis of Variances

## Declarations

The authors declare that they have no conflicts of interest.

## Authors’ Contributions

Pranav Gopalakrishnan: methodology, formal analysis, investigation, data curation, writing - original draft, visualization, and writing - review & editing.

Ahmad Faryami: conceptualization, methodology, formal analysis, investigation, data curation, project administration, visualization, and writing - review & editing.

Carolyn A. Harris: conceptualization, methodology, investigation, resources, supervision, project administration, funding acquisition, and writing - review & editing.

## Acknowledgments

We would like to thank Prashant Hariharan for his assistance with the WSU biorepository. We would also like to thank Adam Menkara for his assistance in data collection, data presentation, and early conceptualization.

